# Learning to evoke complex motor outputs with spatiotemporal neurostimulation using a hierarchical and adaptive optimization algorithm

**DOI:** 10.1101/662072

**Authors:** Samuel Laferriere, Marco Bonizzato, Numa Dancause, Guillaume Lajoie

## Abstract

The development of neurostimulation techniques to evoke motor patterns is an active area of research. It serves as a crucial experimental tool to probe computation in neural circuits, and it has applications in neuroprostheses used to aid recovery of function after stroke or injury. There are two important challenges when designing algorithms to unveil and control neurostimulation-to-motor mappings, thereby linking spatiotemporal patterns of neural stimulation to muscle activation: (1) the exploration of motor maps needs to be fast and efficient (exhaustive search is to be avoided for clinical and experimental reasons) (2) online learning needs to be flexible enough to track ongoing changes in these maps. We propose a stimulation search algorithm to address these issues, and demonstrate its efficacy with experiments in non-human primate models. Our solution is a novel iterative process using Bayesian Optimization via Gaussian Processes on increasingly complex signal spaces. We show that our algorithm can successfully and rapidly learn mappings between complex stimulation patterns and evoked muscle activation patterns, where standard approaches fail. Importantly, we uncover nonlinear circuit-level computations in M1 that would not have been possible to identify using conventional mapping techniques.

## Introduction

Each year, over 15 million people worldwide suffer major debilitating motor system injuries such as spinal cord trauma (*Spinal cord injury*, 2013) or stroke (Thrift et al., 2016). A promising approach to help restore movement applies targeted, artificial stimulation of motor-related neural pathways, e.g. in motor cortex (Cioni B, 2016), spinal cord (Wenger et al., 2016), or peripheral functional electrical stimulation (Selfslagh et al., 2019) using brain-computer interfaces (BCI). New implantable devices which are microfabricated with many (*>*32) electrodes hold potential for targeted and specific stimulation, yet existing control algorithms do not fully take advantage of this, generally relying on incomplete and manual mapping, and often single electrode stimulations. Our goal is to extend these control algorithms to multiple electrode stimulations, which we believe will be necessary to elicit complex movement outputs. Effectively searching the space of possible spatiotemporal stimulation patterns however is a complex task because of its combinatorial explosion in size. Exhaustive search is therefore impossible in practice, especially if algorithms are to be used on-line in clinical settings. Moreover, the mappings between stimulation and output are noisy, and may change over time due to plasticity of neural circuits (Kaas, 1991). Any method to identify stimulation protocols must be robust, and flexible enough to track such changes.

We propose a Gaussian Process (GP) based *hierarchical* Bayesian Optimization (BO) approach. This leverages acquired knowledge of muscle responses for single channel stimulations to build priors for *stim-to-EMG* maps for multi-channel stimulation patterns, where only correction terms need to be learned. The advantages of recursively learning correction terms, rather than a complete map, are threefold: **(1)** Convergence to optimal stimulation requires fewer exploratory stimuli than direct optimization on the space of all signals. **(2)** The algorithm can be used online and adapts quickly to changes in neural dynamics. **(3)** Our method precisely learns the nonlinearities introduced by network dynamics, and can track the evolution of population codes throughout recovery, thus allowing a mapping of circuit-level computations.

In this paper, we describe our novel algorithm and demonstrate its efficacy with motocortical neurostimulation experiments in non-human primates where optimal multi-electrode signals are identified to evoke muscle synergies. We discuss future development directions as well as uncovered circuit-level neural mechanisms present in M1.

## Methods

### Neural Stimulation: Setup and Experiment Description

A 96 channel Utah array is implanted in primary motor cortex (M1) of a male adult Cebus Capella, and five electromyogram (EMG) electrodes are inserted in different muscles of the forearm and hand: *flexor carpi ulnaris, extensor digitorum communis, extensor carpi radialis, opponens pollicis and flexor pollicis brevi* (see monkey in Fig. 1a^5^). We stimulate M1 by sending electrical pulse trains through one or many channels, and observe time series of EMG responses (Fig. 1b) that correlate with the animal’s muscle activations. Our goal is to find the stimulation pattern which will evoke a given target EMG response as best as possible. The Monkey was group housed and supplied with food and water *ad libitum*. The experimental protocol followed the guidelines of the Canadian Council on Animal Care and was approved by the Comité de Déontologie de l’Expérimentation sur les Animaux of the Université de Montréal.

**Figure 1:**
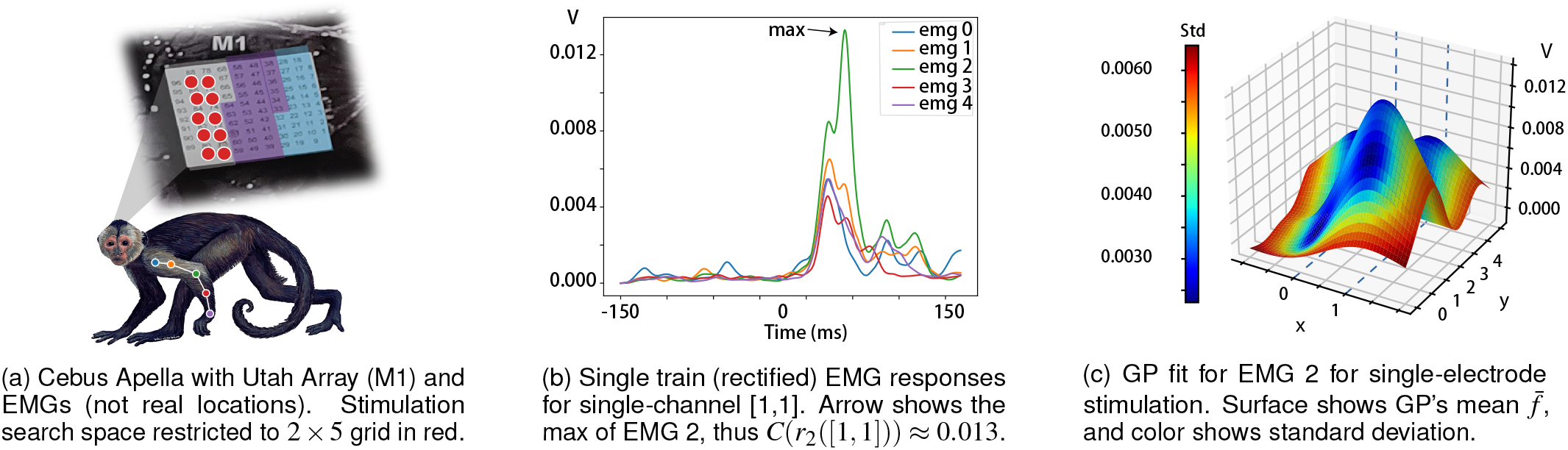
Experiment Setup and Data Description.

Mathematically, we consider electrical stimulation signals that are composed of discrete events (single electrical pulses or short pulse trains) that we call *(stimulation) events* for generality. For our cortical stimulation experiment, we consider a two dimensional 2 × 5 grid that is circled in red in Fig. 1a and use stimulation events consisting of trains of 13 pulses of 30*µ*A, delivered at 330Hz (for a total of roughly 40 ms). Note the spatial configuration of channels is important for learning as Gaussian Processes make use of this information in their kernel distance function. In this article, we restrict our data collection to this small subgrid to allow extensive search of the two electrode stimulation space^6^ (formally defined below). Nevertheless, we note that our algorithm scales nicely to larger channel counts.

We denote each channel by discrete Cartesian coordinates *c* = [*x, y*], *x ∈* {0, 1}, *y ∈* {0,…, 4}. A stimulation containing *k* events is a tuple *s*_*k*_ = (*c*_1_, … *c*_*k*_, ∆*t*_1_, …, ∆*t*_*k*−1_) where *c*_*i*_ indicates the channel of the *i*^th^ event, and ∆*t*_*i*_ is the inter-event interval between events *i* and *i* + 1.^7^ Each *s*_*k*_ generates a noisy response pattern *r*(*s*_*k*_). In our case, we consider the (rectified) response of five EMGs: *r*(*s*_*k*_) = (*r*_0_(*s*_*k*_), …, *r*_4_(*s*_*k*_)). Our goal is to optimize an objective function *C*(*r*(*s*_*k*_)). Here *C* is flexible; it can be extracting the maximum output of a single EMG (or combinations of EMGs), or measuring a distance between evoked pattern *r*(*s*_*k*_) and a target pattern *r*_*target*_. In this paper, *C*(*r*(*s*_*k*_)) returns the maximum output of a single *r*_*i*_(*s*_*k*_) or combinations of *r*_*i*_ (*s*_*k*_) for muscle synergies (see Results), in 150ms following the first stimulus delivery. In general, *r*(*s*_*k*_) could also depend on time. We omit this for notation clarity. We want to find

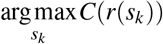

where the argmax can be replaced by argmin if we want to minimize a distance function instead of maximizing an amplitude.

In this article, we demonstrate and test our algorithm offline on the space of double-event stimulations *s*_2_. To do so, we gathered an exhaustive dataset consisting of 10 trials per stimulation pattern (pairs of electrode) for each of these ∆*t*: 0, 10, 20, 40, 60, 80, 100 ms, and sample from this data set to simulate online optimization. An online demonstration will be presented in a forthcoming publication.

### Gaussian Processes for Bayesian Optimization

Given the constraints of our problem, namely that of blackbox derivative-free global optimization under time constraints, Bayesian Optimization (Brochu et al., 2010) is a natural fit. This provides uncertainty estimates that allow tracking and adapting online to both signal delivery changes in the implant and structural changes in the underlying brain substrate.

BO constructs, at every iteration, a probabilistic surrogate to the function *C* being optimized, which is used to balance exploration and exploitation through the design of an acquisition function. It does so by treating the unknown function *C* as a random function and placing a prior over it. This prior dictates attributes of the function such as smoothness and frequency of oscillation. By conditioning on the so far observed responses of the function, a posterior distribution over possible functions is obtained, from which the algorithm can decide where to query next based on optimizing an acquisition function. Acquisition functions convert a probabilistic belief into a deterministic function that explicitly embodies the trade-off between exploration and exploitation. Following the current literature, we choose to model the random surrogate as a Gaussian Process (Rasmussen & Williams, 2005), and use the *Upper Confidence Bound* (Brochu et al., 2010) as acquisition function.

#### Gaussian Process Prediction

GPs are such that for a finite number of training data points ***x*** and their associated responses ***y***, plus a finite number of test data points ***x***_∗_ whose response ***f*** _∗_ we would like to predict, we get a Multivariate Gaussian

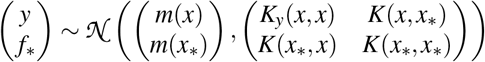

where *m* and *K* are the mean and kernel functions associated to the GP, and 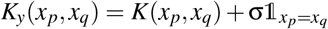, where σ is the noise standard deviation parameter, which will be optimized along with *K*’s parameters. We can get our prediction for ***f***_∗_ by simple conditioning on this Multivariate Normal distribution (Bishop, 2006):

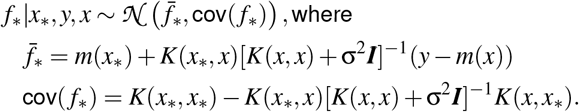

### Sequential Optimization

Given a GP and a set of possible points to explore, we use the Upper Confidence Bound (UCB) acquisition function (Brochu et al., 2010): UCB(*x*) = *µ*(*x*) + *k*σ(*x*) to identify the next query likely to maximize the objective. Notice that *k* is the parameter which modulates the trade-off between exploration (high *k*) and exploitation (low *k*). We set *k* = 3 for all of our experiments, which worked best, but note that the performance of the algorithm is sensitive to this hyperparameter and tuning it by cross-validation or some other method will be necessary for portability across different kinds of neural interfaces.

### Hierarchical GP

We describe here the main (algorithmic) contribution of this paper. Our goal, as previously expounded, is to find the multi-electrode stimulation pattern with the best response, where best is defined by the objective function. The challenge here is that the space of spatiotemporal multielectrode patterns grows combinatorially fast in the number of channels and stimulation event. For two-electrode stimulation on our 2 × 5 grid for example, there are 100 combinations of channel pairs possible, without even considering different inter-event intervals ∆*t*. The direct approach of training a GP on this space is not scalable, and does not take advantage of prior knowledge of motor circuit coding; namely, that motor outputs of spatiotemporal neural activations can often be decomposed (although not exactly) into individual neural-muscle correspondences (Georgopoulos et al., 1986). We leverage this fact in a hierarchical approach where we use GPs fitted on lower dimensional stimuli spaces, to build priors for GPs in higher dimensional stimuli spaces.

More formally for the two-electrode space *s*_2_ = (*c*_1_, *c*_2_, ∆*t*), if we write the single-electrode GP as *f*_1_(*c*) ~ GP(0, *K*_1_) then our prior on the two-electrode GP will be

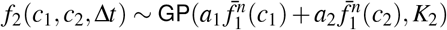

where 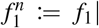 Data is the GP trained on the single-electrode data, 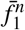 indicates its mean function, and *K*_2_ is a standard Matern52 (Rasmussen & Williams, 2005) multiplicative kernel which separates over time and sapce: 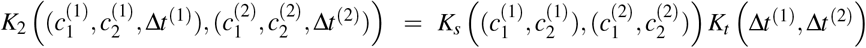

In short, our constructed prior is an independent, additive contribution from the two channels, factoring in the time delay ∆*t*, which is also an explored parameter. We use the kernel in the two-electrode space to learn and correct the multiplicative, nonlinear difference from this prior. The weights *a*_1_ and *a*_2_ and the kernel hyper-parameters are optimized incrementally using BO after each new query. The same procedure can be recursively used to to include more electrodes, although we present results only for the two-electrode case in this paper. We show in the next section that important gains are obtained from using our method.

## Results

### Single Electrode Stimulation

We now describe how we use a GP to build a function approximation of the single electrode responses. In this work, we only use the mean function of this GP as prior information for the two-electrode GP, but we could also incorporate the uncertainty estimates in the two-electrode kernel, or in the likelihood model (see Discussion). Because the ultimate goal is to find the best stimulation in the double electrode space using the minimum amount of queries, we also need to factor in all of the queries made in the single electrode space. For this dataset we find that spending 25 queries in the single electrode space (roughly 2 queries per channel) is enough to have confidently found the highest responding channels, as is illustrated in Fig. 2b.

**Figure 2:**
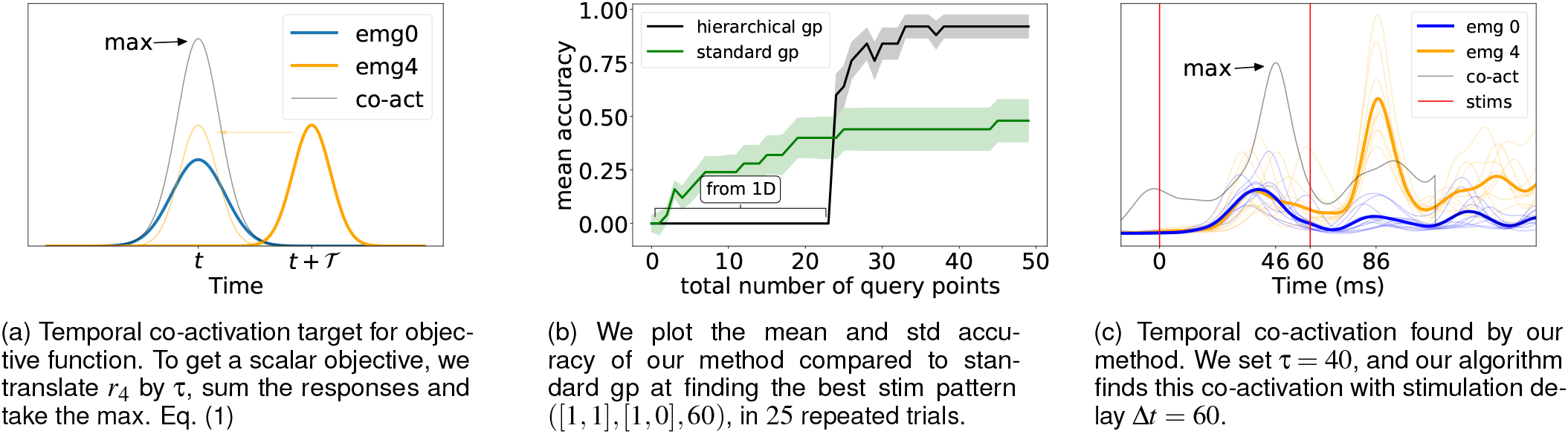
Objective and Results

### Two Electrodes Stimulation Pattern

To showcase our algorithm on a concrete example, we use a temporal co-activation of multiple EMGs (see Fig. 2a) as a target in our objective function, which we define to act as a proxy for a complex sequential two-muscle movement. We target two muscles of the animal, the *flexor carpi ulnaris* (EMG0) and the *opponens pollicis* (EMG4), which we want to activate with a 40ms delay in between peaks. In order to formulate this problem using a similar objective function as described in the Methods section, we make the simplifying assumption that movement amplitude will correlate with the maximum amplitude of combined EMG responses, incorporating the desired delay. We define

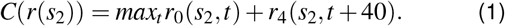

In our search, we restrain the ∆*t* dimension to the discrete set (20, 40, 60)ms due to data collection constraints (see Footnote 6). We found that having the spatial kernel dimensions share lengthscales gave the best results. Furthermore, we constrain this lengthscale to be between 1 and 2 so as to avoid spurious local minima where either the data is explained by noise only (*l* = ∞) or the responses become independent (*l* = 0) (Rasmussen & Williams, 2005). We also constrain the noise standard deviation to be between 5e*−*4 and 1e*−*3 (typical EMG response size are between 1e*−*2 and 3e*−*2), which encompasses the empirical standard deviation of every stimulation pattern. We compare our hierarchical approach, which uses priors built with a GP in the single-electrode space using 20 queries, to a GP which is directly trained on the two-electrode space.

The results in Fig. 2b show that our algorithm clearly outperforms the standard GP-BO procedure, which not only takes much longer to converge, but also is more sensitive to the UCB parameter *k*, and can more easily get stuck in local minima. We show that for the chosen muscle combination, the mean accuracy over 25 repeated trials is 1 after about 40 query points (including the initial ~ 25 queries used to train the GP in the single-electrode space). This means that our algorithm converges to the true best stimulation pattern, namely (*c*1*, c*2, ∆*t*) = ([1, 1], [1, 0], 60), out of a possible 300 (100 channel pairs and 3 time delays ∆*t*) with only 40 total queries. Due to this stimulation pattern having a response much larger than other channels (0.041V, whereas other high-responding patterns are around 0.030V), we believe this metric (which focusses on the very best channel) to be a good measure of success. In other words, any other stimulation pattern would not have resulted in a movement as obvious and well-defined as this one. We show in Fig. 2c the resulting co-activation found.

## Conclusion and Discussion

We showed that the hierarchical approach to build GPs on the space of multi-electrode stimulation is a viable one. Not only does it far outperform the standard GP approach, showing great potential for use in stroke recovery, but it can also be used online to find the optimal stimulation strategy for a desired co-activation output. This is a step forward in linking brain activity and behavior by being able to control neurons directly (Alivisatos et al., 2013).

The most novel and interesting part of our work is the ability to control and elicit complex movements online. We used a restricted stimulation search space for data collection purposes, and clearly demonstrated faster learning using our approach. Future work will focus on testing this algorithm online, using all of the 96 electrodes in the Utah array of our example setup, which will require a stopping criterion for the single electrode space search.

We made a few simplifying modeling assumptions, which could easily be improved upon, though we are unsure whether this will be necessary. For one, we assumed homoscedastic noise, whereas biology is better described by Poisson noise. Second, we trained the different EMG GPs (for each *r*_*i*_) independently, whereas they are clearly correlated (see Fig. 1b), and could share information through multi-output (also called co-Kriging) models. Thirdly, using a non-stationary kernel could accelerate the search even more. This would allow having a large lengthscale for most of the search space, yet have a smaller lengthscale near optimal stimulation patterns to permit finding the true maximum.

Finally, we note that our method can easily be adapted to more complex objective functions such as incorporating both forelimb and backlimb movements, and to different sensor modalities such as accelerometer data or 3D position using cameras.

## Acknowledgments

We thank Maximilian Puelma Touzel and Olivier Caron-Grenier for useful discussions. Funding: MB [IVADO fellwoship], ND [FRQNT group grant (2019-PR-253402)], GL [NSERC Discovery Grant (RGPIN-2018-04821), FRQNT Young Investigator grant (2019-NC-253251), FRQS Research Scholar Award, Junior 1 (LAJGU0401-253188)]

## Footnotes

5 Royalty-free image from unixtitan.net

6 With this current setup, the data collection protocol takes about an hour, during which the monkey needs to receive intravenous ketamine delivery every 8 minutes. It would be hard to maintain the animal in an overall stable state for longer times, and this could change the data distribution

7 In this experiment, we use trains of fixed intensity but power could be added to the stimulation parameters with the same formalism

